# Sequencing of small RNAs of the fern *Pleopeltis minima* (Polypodiaceae) offers insight into the evolution of the microRNA repertoire in land plants

**DOI:** 10.1101/078907

**Authors:** Florencia Berruezo, Flavio S. J. de Souza, Pablo I. Picca, Sergio I. Nemirovsky, Leandro Martinez-Tosar, Mercedes Rivero, Alejandro N Mentaberry, Alicia M. Zelada

## Abstract

MicroRNAs (miRNAs) are short, single stranded RNA molecules that regulate the stability and translation of messenger RNAs in diverse eukaryotic groups. Several miRNA genes are of ancient origin and have been maintained in the genomes of animal and plant taxa for hundreds of millions of years, and functional studies indicate that ancient miRNAs play key roles in development and physiology. In the last decade, genome and small RNA (sRNA) sequencing of several plant species have helped unveil the evolutionary history of land plant miRNAs. Land plants are divided into bryophytes (liverworts, mosses), lycopods (clubmosses and spikemosses), monilophytes (ferns and horsetails), gymnosperms (cycads, conifers and allies) and angiosperms (flowering plants). Among these, the fern group occupies a key phylogenetic position, since it represents the closest extant cousin taxon of seed plants, i.e. gymno- and angiosperms. However, in spite of their evolutionary, economic and ecological importance, no fern genome has been sequenced yet and few genomic resources are available for this group. Here, we sequenced the small RNA fraction of an epiphytic South American fern, *Pleopeltis minima* (Polypodiaceae), and compared it to plant miRNA databases, allowing for the identification of miRNA families that are shared by all land plants, shared by all vascular plants (tracheophytes) or shared by euphyllophytes (ferns and seed plants) only. Using the recently described transcriptome of another fern, *Lygodium japonicum*, we also estimated the degree of conservation of fern miRNA targets in relation to other plant groups. Our results pinpoint the origin of several miRNA families in the land plant evolutionary tree with more precision and are a resource for future genomic and functional studies of fern miRNAs.

## Introduction

Land plants (embryophytes) evolved from fresh water, streptophyte green algae during the Ordovician period, over 470 million years ago (MYA) [1, 2]. Major diversification events between the Silurian and Permian periods (385-470 MYA) gave rise to groups represented today by bryophytes *sensu lato* (liverworts, hornworts and mosses), lycophytes (clubmosses and spikemosses) and the euphyllophytes, comprising monilophytes (ferns) and spermatophytes (seed plants) [1]. Extant seed plants are divided into gymnosperms (cycads, conifers) and angiosperms. Early diversification in the Palaeozoic resulted in different strategies of alternating (gametophytic and sporophytic) generations and the origination of key anatomical features in different groups of plants, including vascular systems and various types of meristems, leaves and roots, as well as seeds [1, 3]. The origin and evolution of these and other features is studied with fossils and comparative anatomical and physiological studies, but comparative genomics of extant species can also help understand plant evolution. In the last decade, several angiosperm genomes have been sequenced, as well as the genomes of a moss (*Physcomitrella patens*), a lycopod (the spikemoss *Selaginella moellendorffii*) and a conifer tree (Norway spruce, Picea abies) [4-6]. Genomic analyses show that land plants share a common set of transcriptional regulators, including transcription factors and microRNAs, that set them aside from green algae [3, 7].

MiRNA genes are transcribed as long, primary RNA precursors (pri-miRNAs) that can form a foldback, hairpin structure. In plants, pri-mRNAs are processed by Dicer-like 1 (DCL1) protein to generate a miRNA/miRNA* duplex. The miRNA* ("star") strand is generally degraded, whereas the mature miRNA molecule is incorporated into a RNA-induced silencing complex (RISC) that contains Argonaute 1 protein (AGO1). The ~21nucleotide (nt) miRNA associated with RISC serves as a guide to recognize specific messenger RNAs (mRNAs), interfering with their translation or leading to mRNA degradation [8, 9]. MiRNAs are, thus, negative regulators of gene expression. Analyses of land plant genomes and small-RNA transcriptomics have revealed that many miRNA genes are common to all plant groups, suggesting that they originated before or during the colonization of land by plant ancestors. Many of these deeply conserved miRNAs play important roles in plant development and have likely contributed to the diversification of land plant body plans during evolution [7, 10-13].

Our understanding of plant miRNA evolution is hampered by biased sampling, since most genomic and transcriptomic studies have been carried out in angiosperms. In particular, there is a dearth of knowledge on the miRNA repertoire of ferns. The fern clade (Monilophyta) [14] comprises today around 12,000 species. It occupies a special place in the plant phylogenetic tree as the sister group of seed plants (Fig 1A). Thus, the study of ferns should illuminate critical aspects of plant evolution, like the transition from homospory to strict heterospory, sporophyte dominance, as well as aspects of the development of leaves, roots and vascular systems that distinguish ferns and seed plants [3, 15]. No fern nuclear genome has been sequenced to date, although some genomic surveys and transcriptomic analyses have been carried out [16, 17]. An early survey based on hybridization arrays indicated the presence of eight miRNAs in the fern *Ceratopteris thalictroides* that are also conserved in other land plants [18]. More recently, a broad survey of several land plants included an aquatic fern, *Marsilea quadrifolia*, which added some conserved miRNAs to the fern repertoire [19].

**Fig. 1.**
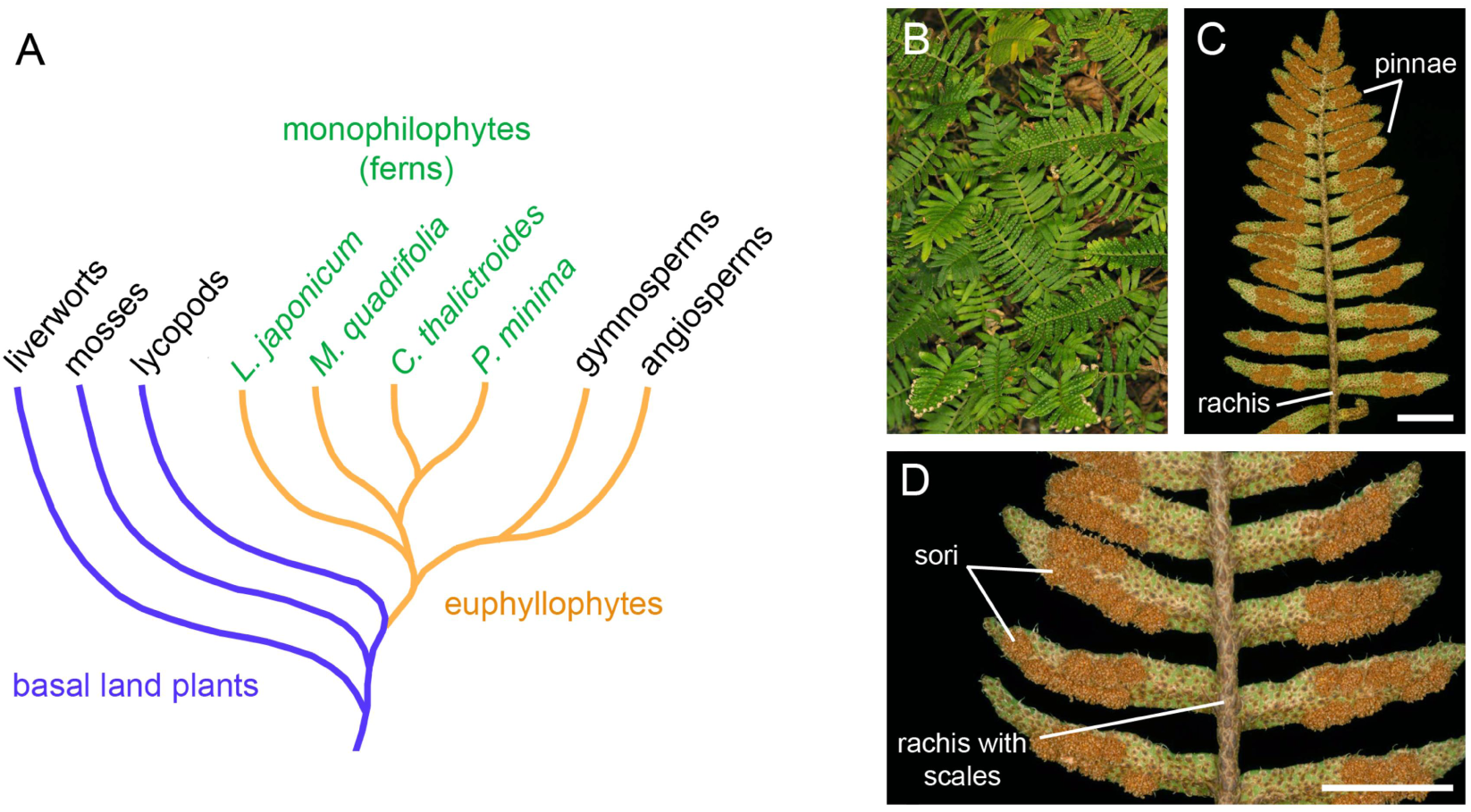
Phylogenetic position and habit of *Pleopeltis minima*. **(A)** Simplified summary of phylogenetic relationships among major plant lineages relevant for the present study [3, 14, 29]. The fern (“monilophytes”) clade is the sister group to seed plants, gymnosperms and angiosperms. Within ferns, *P. minima* belongs to the order Polypodiales and is more closely related to *Ceratopteris thalictroides*, while *Marsilea quadrifolia* belongs to the order Salviniales and *Lygodium japonicum* to the order Schizaeales. **(B)** Habit of *P. minima* growing as epiphyte upon a tree trunk. **(C)** Abaxial side of leaf surface showing form and organization. **(D)** Portion of fertile leaf showing position of sori and scales. Scale bars: 5 mm.

Here, we study a South American fern, *Pleopeltis minima* (Bory) J. Prado & R.Y. Hirai (synonyms: *Polypodium squalidum* Vell.; *Pleopeltis squalida* (Vell.) de la Sota) [20]. *P. minima* is a small, mostly epiphytic fern belonging to the Polypodiaceae family that occurs in forests in Argentina, Bolivia, Southern Brazil, Paraguay and Uruguay. Its scaly rhizome bears several 2-10 cm long compound leaves (fronds), divided into pinnae (leaflets). Fertile leaves possess exindusiate sori (cluster of sporangia) of round shape on the abaxial side (Fig 1B). One particular characteristic of *P. minima* is its dissecation tolerance (poikilohydry) [21, 22]. We have sequenced the small RNA fraction of *P. minima* and searched for miRNAs phylogenetically conserved in other plants. We present a group of conserved fern miRNAs and their predicted targets, thereby advancing our understanding of miRNA evolution in the fern and land plant lineage.

## Results

### Sequencing and analysis of P. minimas RNAs

Small RNAs (sRNAs) were extracted from two samples of fertile fronds of *P. minima*, converted to cDNA and subjected to Illumina sequencing, generating a total of 25,947,729 reads, of which 20,989,537 were high-quality reads with lengths between 15 and 45 nt. Of these, 6,268,973 reads corresponding to mRNA fragments, tRNAs, rRNAs, snoRNAs, snRNAs and repetitive elements were discarded, leaving a total of 14,720,564 sRNA reads to be further analysed (S1 Table).

The length of functional sRNAs are thought to lay in the 20-24 nucleotide (nt) range. In angiosperms, sRNA length distribution typically displays two peaks at 21 and 24 nt, with miRNAs centred around 21 nt, while 24 nt sRNAs are associated with silencing of retroposons by heterochromatinization [9]. The lycopod *S. moellendorffii* and the spruce *Picea abies* both lack a prominent 24 nt sRNA peak [5, 6]. In *P. minima*, however, we observed a prominent 21 nt peak and a smaller, but noticeable, peak at 24 nt (Fig 2). A similar distribution is found in the aquatic fern *M. quadrifolia* [19], showing that the formation of sRNAs of both classes is a common characteristic of ferns.

**Fig. 2.**
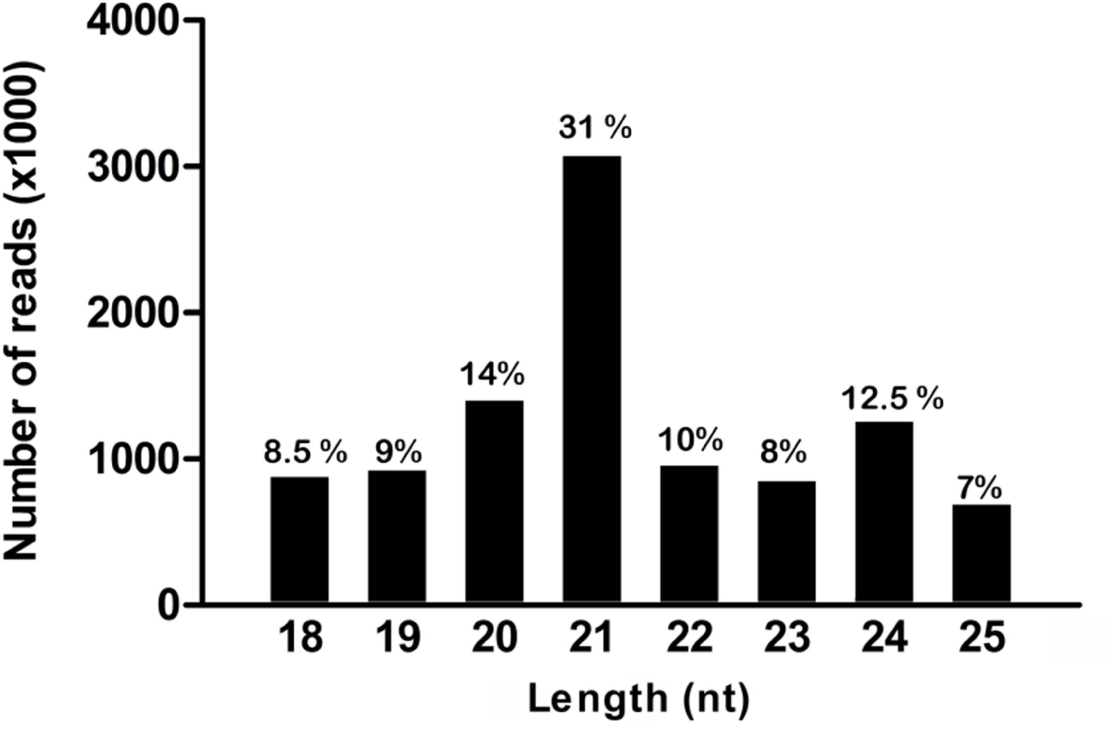
Size distribution of P. minima sRNA sequences. The graph displays the raw number of sRNA reads in the range of 18 to 25 nt. The percentage of reads in each size category in the range is indicated. A large peak at 21 nt and a smaller peak at 24 nt are noticeable.

### Identification of conserved *P. minima* miRNAs

Since *P. minima* lacks a genome sequence and no fern genome is available for comparison, we focused on finding miRNAs in our samples that are conserved in other land plant genomes. To this end, the 16,482,278 sRNA reads of *P. minima* were compared to version 21 of miRBase [23], yielding 1,127,970 reads that could be aligned to miRNAs or MIRNA genes in the database, corresponding to 3,912 unique sequences (S1 Table).

It is known that the confidence on the existence of miRNA families in miRBase varies widely, and the evidence for many miRNA families is rather weak, specially for miRNAs coming from species lacking a genome sequence [8]. In view of this, to identify bona fide *P. minima* miRNAs we required the miRNA families 1) to be represented by at least 10 reads of 20-22 nt in our data; 2) to exhibit consistent 5’ cleavage processing of the mature miRNA strand and 3) to be identical or very similar to highly confident sequences of mature miRNA or miRNA* strands of other plant species, as defined by [8]. For some miRNA families, namely miR162, miR395 and miR477, even though the total number of reads was under 10, we still considered them to be real miRNAs since the reads included both miRNA and miRNA* strands that were identical or nearly identical in sequence to the corresponding miRNA and miRNA* sequences from other plants (S1 Fig).

The analysis revealed a total of 57 conserved miRNAs in *P. minima*, belonging to 23 miRNA families (Table 1). Since miR156 is related in sequence and function to miR529 [24, 25] and the same occurs between miR159 and miR319 [26, 27], the number of conserved *P. minima* miRNA families could alternatively be considered as being 21. For five families (pmi-miR162, pmi-miR390, pmi-miR395, pmi-miR477 and pmi-miR529), sequences that represent the putative miRNA/miRNA* duplex strands (i.e., the 5p and 3p strands of the miRNAs) were identified (S1 Fig). Sixty-one percent (35/57) of the conserved *P. minima* miRNAs, belonging to 18 of the 23 miRNA families, have a 5’ terminal uridine residue (Table 1), a conserved feature of miRNAs recognized by the AGO1 protein, while miRNAs from the pmi-miR390, pmi-477 and pmi-miR529 families have mostly a 5’ terminal adenine residue, a feature found in miRNAs recognized preferentially by AGO2 and AGO4 in angiosperms [28].

**Table 1.**
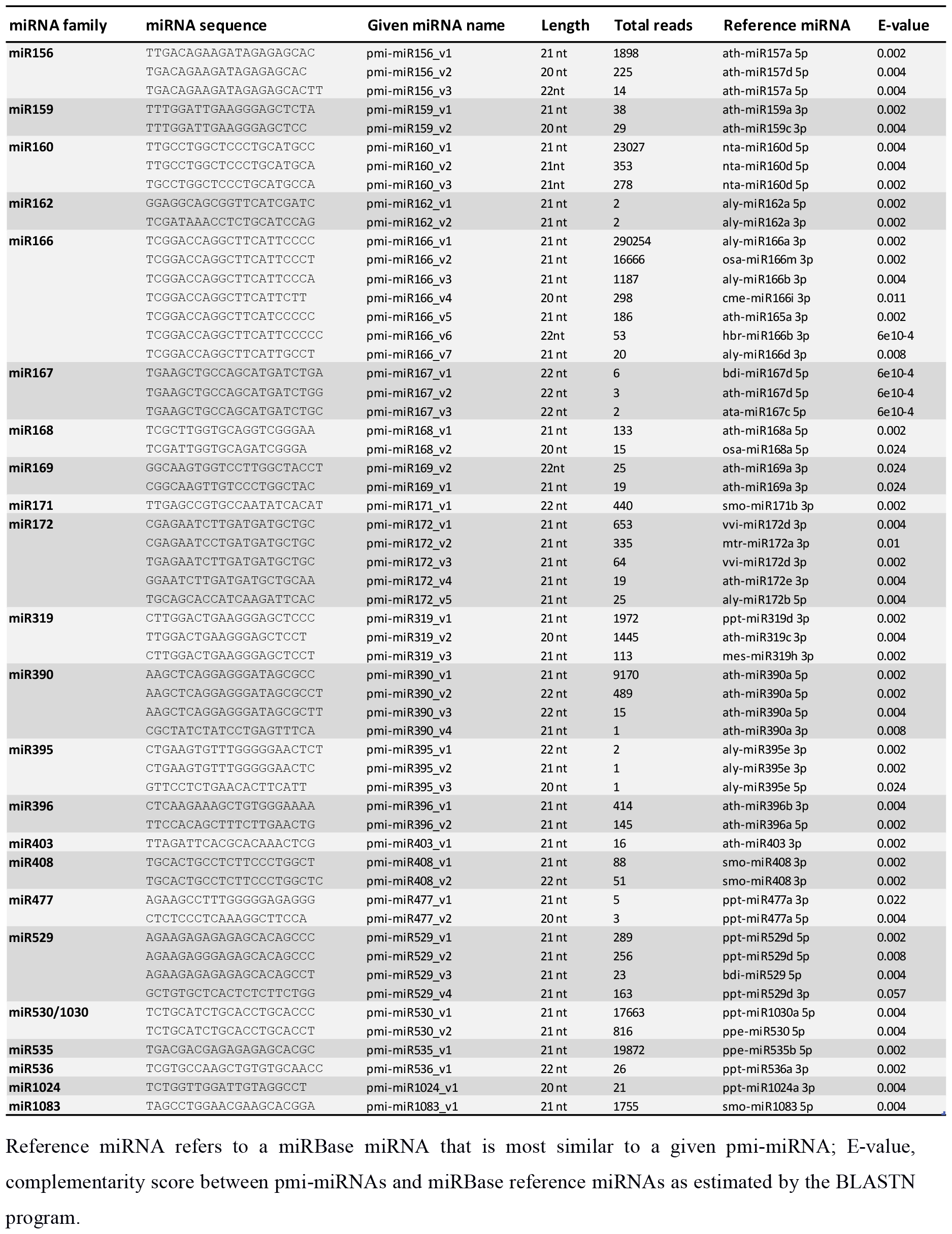
Conserved *P. minima* miRNAs.

### Ancient land plant miRNA families

MiRBase contains miRNAs from well-characterized plant species for which genome information is available, like the moss P. patens, the lycopod S. moellendorfii and the angiosperms Arabidopsis thaliana and Oryza sativa [8, 10], but miRNAs coming from many recent sequencing surveys are not present in the database. To expand the scope of our phylogenetic analysis, we compared the conserved *P. minima* miRNAs to those of *Marsilea quadrifolia* [19], a water fern that diverged from polypod ferns like *P. minima* around 220 MYA [14, 29]. The miRNAs of *M. quadrifolia* and seed plants identified by Chávez-Montes et al. [19] were reanalysed to identify bona fide conserved miRNAs (see Materials and Methods). We also compared *P. minima* sequences with the miRNAs of various gymnosperms and basal angiosperms [19, 30-33], as well as conserved miRNAs identified recently in the liverworts *Pellia endiviifolia* [34] and *Marchantia polymorpha* [35, 36]. This allowed us to trace the history of miRNA origin and conservation in land plants, with emphasis on the fern miRNA repertoire.

Particularly noteworthy are miRNA families miR156/319, miR160, miR166, miR171 and miR408, which are found in all land plant groups, including liverworts, which represent the earliest-branching group in land plant evolution (Figs 1A and Fig 3). Some ancestral miRNAs have a more patchy phylogenetic distribution and are missing in a few basal groups. For instance, miR396 is absent from moss, miR167 and miR530 are missing from liverworts and lycopod, while miR535 and miR390 are specifically missing in the lycopod. In any case, the *P. minima* miRNA repertoire is conserved compared to that of basal land plant groups. Thus, all 12 ancient miRNAs present in liverworts (P. endiviifolia and/or M. polymorpha) and all 15 ancient miRNAs of the moss, are present in *P. minima*. Similarly, all 10 ancient miRNAs found in the lycopod are also present in *P. minima* (Fig 3).

**Fig. 3.**
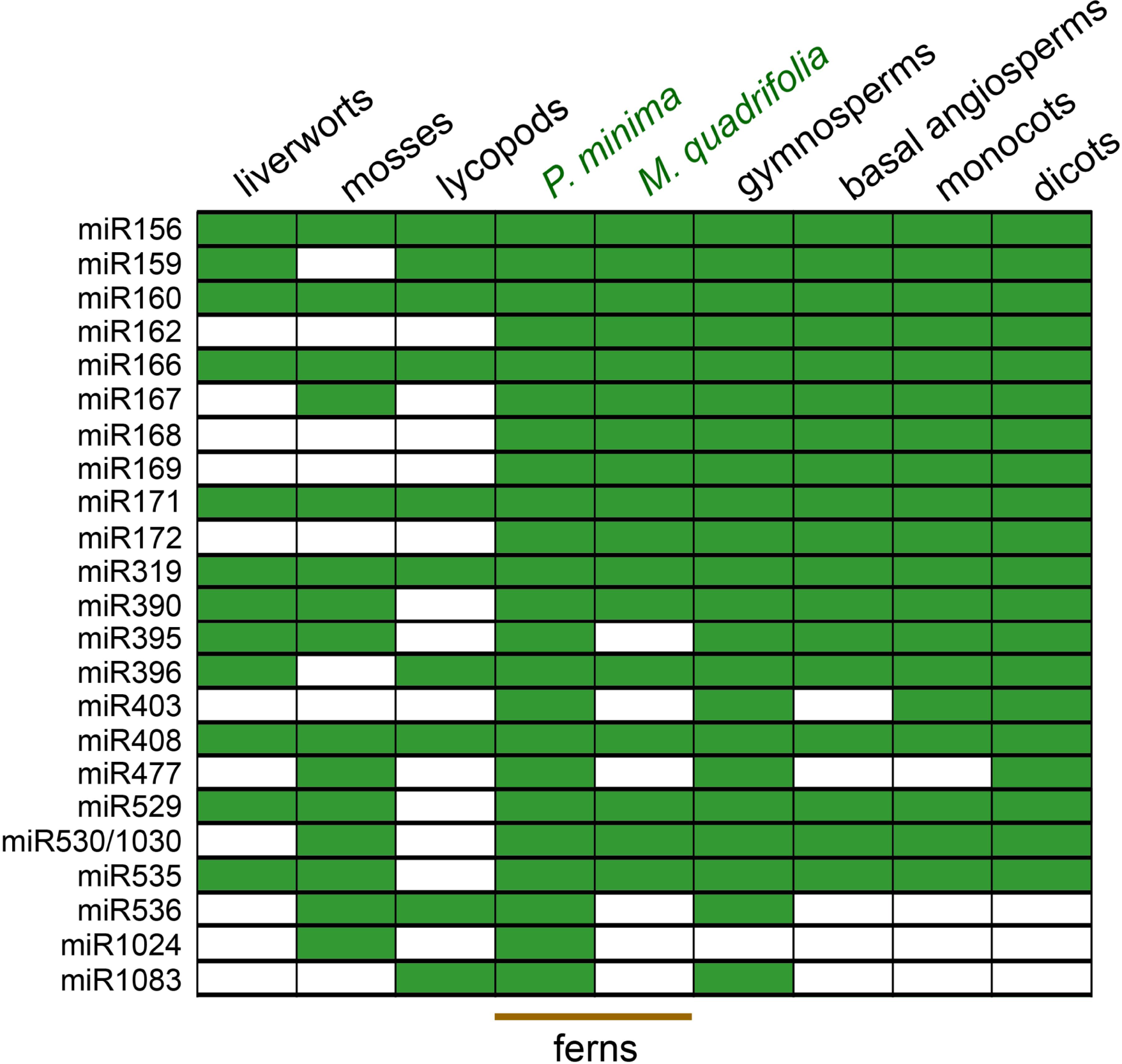
Phylogenetic distribution of conserved fern miRNAs. The grid indicates the presence (green) or absence (white) of miRNA families in liverworts (*M. polymorpha and/or P.endiviifolia*), mosses (*P. patens*), lycopods (*S. moellendorffii*), ferns (*P. minima* and *M. quadrifolia*) as well as gymnosperms, basal angiosperms, monocots and dicots.

Some miRNA families are only present in a few basal groups. One is miR1024, a high-confidence miRNA from the moss P. patens [10] which is found in *P. minima* and in no other species, not even *M. quadrifolia* (Fig 3). Although this miRNA has only 21 reads in our samples, it displays 100% identity over 20 nt to P. patens miR1024, indicating that it is a real miRNA. Thus, it might represent an ancient miRNA that has been independently lost from several lineages, including seed plants. Another case is that of miR1083, which is found in *P. minima* (but not in *M. quadrifolia*, see Materials and Methods) with over 1,700 reads (Table 1). This miRNA is present in the lycopod, *S. moellendorffii* [10], is abundant in gymnosperms [19, 31, 33], but cannot be identified in angiosperms (Fig 3; see Materials and Methods). Thus, miR1083 seems to be an ancient tracheophyte miRNA that has been specifically lost from angiosperms.

### Euphyllophyte miRNAs

We found that some miRNAs families are only common to ferns (*P. minima*, *M. quadrifolia* or both) and seed plants (gymnosperms and angiosperms), suggesting that these miRNAs originated in the early evolution of euphyllophytes. These are miR162, miR168, miR169, miR172 and miR403 (Fig 3). Among these euphyllophyte miRNAs, the case of miR403 is noteworthy since this miRNA cannot be detected in *M. quadrifolia* nor in most non-dicot groups [19, 37] (see Materials and Methods). However, miR403 has recently been identified in gymnosperms like the Chinese fir, *Cunninghamia lanceolata* [30] and the spruce, *Picea abies* [33], suggesting a complex evolutionary history for this miRNA family.

### Conserved fern miRNA targets

Extensive experimental evidence from several angiosperms and the moss *P. patens*, have unveiled that the mRNAs targeted by ancestral miRNAs are often also deeply conserved [7, 10, 11, 38]. Recently, degradome data has shown that the targets of liverwort ancestral miRNAs are also partially conserved with that of other lineages [34-36]. As for ferns, the only available information consists of putative mRNA targets for *C. thalictroides* miR160, miR171 and miR172, which were also found to be conserved with those of other plant lineages [18].

To confirm and expand the evidence for conservation of ancestral miRNA targets in ferns, we computationally predicted the mRNA targets of conserved *P. minima* miRNAs. Since no *P. minima* transcriptome is available, we made use of the extensive transcriptome data obtained for the fern *Lygodium japonicum* [17]. *L. japonicum* belongs to the Schizaeales order, which places it as a distant cousin to polypod ferns like *P. minima* [14, 29]. We compared the conserved *P. minima* miRNA repertoire to the *L. japonicum* transcriptome using the psRNATarget platform [39] and searched for putative targets that had previously been found in other land plants.

The analysis (Table 2) indicates that 11 out of the 23 *P. minima* miRNA families have the potential to target *L. japonicum* RNAs that are also targeted in other species. Thus, pmi-miR156, as well as pmi-miR529, both target a mRNA that encodes a Squamosa Promoter Binding (SPB) transcription factor; pmi-miR159 and pmi-miR319 both target mRNAs that encode transcription factors of the MYB family; pmi-miR160 targets mRNAs encoding transcription factors of the Auxin Response Factor (ARF) family; pmi-miR166 targets a mRNA encoding a transcription factor of the class III homeodomain-leucine zipper (Class III HD-Zip) family; pmi-miR171 targets a GRAS-domain transcription factor mRNA; miR172 targets the mRNA of a Apetala-2 transcription factor (Table 2). Targets homologous to these have been observed in experimental work done in angiosperms, moss, or both [10, 11, 38]. The *P. minima* miR160, miR171 and miR172 predicted targets confirm those of Axtell and Bartel [18] for *C. thalictroides*. Recently, a thorough survey found that ferns possess three clades of Class III HD-Zip genes [40], and we found that pmi-miR166 can potentially target all Class III HD-Zip paralogues of the fern Psilotum nudum identified in that study (S1 Fig).

**Table 2.** Putative conserved fern targets of *P. minima* miRNAs.

| miRNA family | *Lygodium* transcript | E | Transcript function | Target conservation |
| --- | --- | --- | --- | --- |
| pmi-miR156 | Locus_14997 | 1 | SBP transcription factor | liverworts, moss, gymnosperms, angiosperms |
| pmi-miR159 | Locus_7173 | 1.5 | MYB transcription factor | gymnosperms, angiosperms |
|  | Locus_3676 | 2 | RWP-RK domain transcription factor | same target as miR319 in liverworts |
| pmi-miR160 | Locus_6056 | 0 | Auxin response factor (ARF)-B3 DNA binding domain | liverworts, moss, *Ceratopteris*, gymnosperms, angiosperms |
|  | Locus_3364 | 1 | Auxin response factor (ARF)-B3 DNA binding domain | liverworts, moss, *Ceratopteris*, gymnosperms, angiosperms |
| pmi-miR166 | Locus_1230 | 3.0 | Class III homeodomain-leucine zipper protein C3HDZ2 | liverworts, moss, gymnosperms, angiosperms |
| pmi-miR168 | Locus_158 | 4.0 | Argonaute 1 (AGO1) | gymnosperms, angiosperms |
| pmi-miR171 | Locus_11016 | 1.5 | GRAS domain transcription factor (SCL6) | liverwort (*Marchantia*), moss, *Ceratopteris*, gymnosperms, angiosperms |
| pmi-miR172 | Locus_9880 | 0.5 | Apetala2 (AP2) | *Ceratopteris*, gymnosperms, angiosperms |
|  | Locus_1423 | 1 | Apetala2 (AP2) | *Ceratopteris*, gymnosperms, angiosperms |
| pmi-miR319 | Locus_7173 | 2.0 | MYB transcription factor | liverwort (*Marchantia*), moss, gymnosperms, angiosperms |
|  | Locus_831 | 3.0 | MYB transcription factor | liverwort (*Marchantia*), moss, gymnosperms, angiosperms |
|  | Locus_1866 | 3.0 | MYB transcription factor | liverwort (*Marchantia*), moss, gymnosperms, angiosperms |
|  | Locus_3676 | 2.0 | RWP-RK domain transcription factor | liverworts (*Marchantia*, *Pellia*) |
| pmi-miR390 | Locus_39179 | 0.5 | *TAS3* tasi-RNA targeting ARF3/4 | liverwort (*Marchantia*), moss, gymnosperms, angiosperms |
|  | Locus_20755 | 0.5 | *TAS3* tasi-RNA targeting ARF3/4 | liverwort (*Marchantia*), moss, gymnosperms, angiosperms |
| pmi-miR408 | Locus_5439 | 3.0 | Polyphenol oxydase (PPO) | liverwort (*Pellia*) |
| pmi-miR529 | Locus_14997 | 1 | SBP domain transcription factor | liverwort (*Marchantia*), gymnosperms, angiosperms |

E-complementarity score between miRNA and target RNA as estimated by the psRNATarget program

The miR390 family does not control protein-coding mRNAs, but rather targets a group of non-coding transcripts called TAS3 RNAs (trans-acting siRNAs). In the moss *P. patens* and in angiosperms, two miR390 molecules hybridize to different segments of a *TAS3* RNA, which is then processed by a complex mechanism to produce one or more 21-nt ta-siRNA molecules that are incorporated into a RISC complex that targets the mRNAs of *ARF3* and *ARF4* transcription factors [41]. In addition, *P. patens TAS3* can also produce ta-siRNAs that target the transcription factor AP2 [10], and there is evidence that the tasiRNAs generated from the *TAS3* of the liverwort *M. polymorpha* can target *AP2* as well [36, 42]. As for the *P. minima* miR390 family, we found that they can potentially target two *L. japonicum* non-coding transcripts which exhibit characteristics of *TAS3* RNAs (Table 2). These *L. japonicum TAS3* transcripts carry two sites predicted to hybridize to pmi-miR390; in addition, between the miR390 sites one potential ta-siRNA, derived from the (+) strand of the RNA, can be produced that could target *L. japonicum* ARF3/4 mRNAs (S2 Fig), indicating that the control of ARF genes by miR390 is conserved in ferns.

Of particular interest are some predicted *P. minima* miRNA targets that are conserved with liverwort miRNAs but not other plants. Specifically, apart from transcription factors of the MYB family, pmi-miR159 and pmi-miR319 potentially target a mRNA that encode a RWP-RK domain-containing protein (Table 2). Degradome studies in the liverworts *P. endiviifolia* and *M. polymorpha* identified mRNAs encoding RWP-RK proteins as bona fide targets of miR159 and miR319 homologues of these species [34, 35]. Interestingly, the pairing between pmi-miR159/319 and its potential *L. japonicum* RWP-RK target occurs in the 3’-UTR of the mRNA, the same region that is targeted in the homologous liverwort *M. polymorpha* mRNA by miR319 (S3 Fig). Since mRNAs encoding RWP-RK proteins have not been identified as targets of miR159/319 in moss, gymnosperms or angiosperms [10, 11, 30, 31, 33, 38, 43], it is possible that the miR159/319-mediated regulation of RWP-RK proteins is an ancient land plant feature that has been lost from other lineages. Another target possibly conserved between ferns and liverworts is the mRNA encoding the enzyme polyphenol oxydase (PPO), which was identified as a potential target of pmi-miR408 (Table 2). In the liverwort *P. endiviifolia*, *PPO* was also identified as an experimental target of miR408 by degradome analysis [34]. Significantly, miR408 targets the 3’-UTR of *PPO* mRNAs in both ferns and liverworts (S4 Fig) [34].

In seed plants, some miRNAs target genes involved in the biogenesis and function of miRNA themselves. Thus, in angiosperms, miR162 targets Dicer-like 1 (DCL1), miR168 targets AGO1 and miR403 targets AGO2 [11]. In our analyses, *P. minima* miR162 and miR403 are not predicted to target genes similar to DCL1 or AGO2, but pmi-miR168 is predicted to target *L. japonicum* AGO1 (Table2). The sequence predicted to be targeted by pmi-miR168 is located within the 5’ half of the *AGO1* coding region, outside the segments that encode the conserved domains of the protein, and corresponds to the homologous *AGO1* region targeted by miR168 in A. thaliana (S5 Fig) [44]. Thus, the negative feedback loop involving miR168 and *AGO1* may be an ancestral euphyllophyte feature.

### Discussion

In this work, we identify the conserved miRNAs of a polypod fern, *P. minima*, by comparing its miRNA repertoire to those of other land plant groups. Figure 4 shows a revised scenario for the emergence of different miRNA families during early land plant evolution. Concerning the fern miRNAome, our analyses allowed us to reach a series of conclusions. First, we found that 16 ancient miRNA families that originated in the beginning of land plant evolution, as evidenced by their presence in liverworts and/or mosses, are all present in *P. minima*, showing that the fern miRNA repertoire is very conservative in this regard (Fig 4). This is not a trivial observation, as illustrated by the lycopod *S. moellendorffii*, which lacks half of the ancestral miRNAs that are present in bryophytes (Fig 3). Second, by comparing *P. minima* miRNAs and those of the aquatic fern, *M. quadrifolia*, with the rest of land plants, we identified five miRNAs that seem to have originated at the beginning of euphyllophyte evolution, before the split of the lineages of ferns and seed plants (gymnosperms and angiosperms; Fig 4). The miRNAome of *P. minima* seems to be more conservative than that of *M. quadrifolia*, since six ancestral miRNAs found in the former could be identified in the latter (Fig. 3). Third, by searching putative targets of *P. minima* miRNAs using the transcriptome of another fern, *L. japonicum*, we provide evidence that the targets of 11 fern miRNA families are deeply conserved in land plant evolution.

**Fig. 4.**
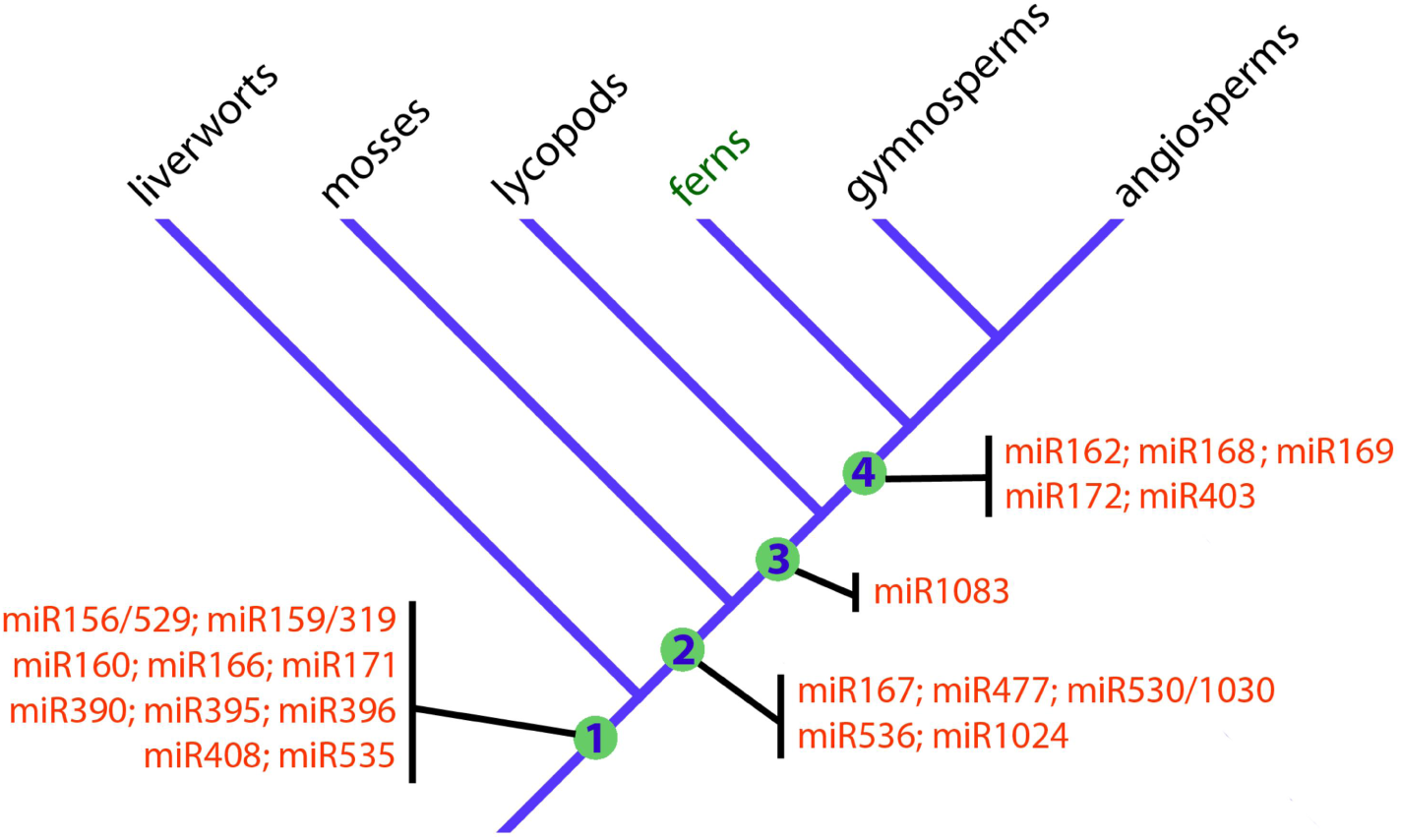
Emergence of miRNAs during early land plant evolution. The schematic cladogram shows the origin of ancient miRNA families during early land plant evolution. Only miRNA families that originated in early land plant evolution are shown, namely land plants (Embryophyta, node 1), mosses and vascular plants (node 2), vascular plants (Tracheophyta, node 3) and ferns and seed plants (Euphyllophyta, node 4). Note that five miRNA families seem to be restricted to euphyllophytes and that only miR1083 seems to have originated in the branch leading to vascular plants.

Molecular studies on basal land plant groups can give crucial clues on the evolution of plant form [3]. As gene expression regulators, miRNAs are essential for proper plant development and may have played important roles in the early evolutionary diversification of land plants, making comparative functional studies of ancestral miRNAs in different plant groups very interesting in this regard [7]. Some ancestral miRNAs present in *P. minima* exhibit an intriguing phylogenetic distribution and are interesting candidates for functional studies. One of these is miR1083, a tracheophyte-specific miRNA that is present in lycopods and is abundantly expressed in *P. minima* and gymnosperms, but is apparently absent from angiosperms (Figs 3 and 4) [19]. The other is miR1024, which is present in moss and *P. minima*, but has not been detected in any other species investigated so far (Fig 3) [10]. Neither the expression pattern nor the targets of miR1083 and miR1024 have been studied in any plant, but their antiquity and conservation make them interesting subjects for future study.

Our bioinformatic search for potential targets of fern miRNAs indicate that 11 of the ancestral pmi-miRNAs might target genes that are homologous to those of other land plants (Table 2). Although our evidence is based on an in *silico* approach, the conservation of potential miRNA/target pairs in species that are separated by hundreds of millions of years of independent evolution is strong evidence that these fern transcripts are real targets. Among these, we found that pmi-miR390 could target *TAS3* RNAs with the potential to generate tasi-RNAs that, in their turn, target genes encoding ARF3/4 transcription factors from *L. japonicum* (S4 Fig). This represents the first full description of *TAS3* transcripts in ferns and an indication that miR390 function is conserved in this group, as suggested by a previous PCR-based survey [42].

Another interesting fern conserved target was found for pmi-miR166, which can target transcription factors of the Class III HD-Zip (C3HDZ) family. In angiosperms, C3HDZ genes are expressed in the adaxial side of developing leaves and are necessary for the proper dorsoventral patterning of these structures [45]. In the abaxial side, on the other hand, C3HDZ expression is repressed by miR166, which is also necessary for proper leaf patterning in angiosperms [46, 47]. Even though it is often considered that the leaves of ferns and seed plants evolved independently [3, 48], Vasco et al. [40] have recently observed that C3HDZ genes are specifically expressed in the adaxial side of developing fern leaves, just like in angiosperms, giving credence to the hypothesis that euphyllophyte megaphylls are homologous structures. Considering our observation that pmi-miR166 could target fern C3HDZ genes, future studies on the expression and function of fern miR166 may help establish the extent of molecular homology in the gene regulatory network that controls leaf development in euphyllophytes.

A recurrent feature in the evolution of plant gene regulation is that certain mRNAs encoding proteins involved in miRNA processing and function are themselves controlled by miRNAs. In angiosperms, *AGO1*, *DCL1* and *AGO2* are targets of miR168, miR162 and miR403, respectively [11], and *AGO1* is also a miR168 target in the Chinese fir, a gymnosperm [30]. Interestingly, liverworts and mosses also target *AGO1* and *DCL1* with a different set of miRNAs. Thus, miR902 and miR1047 target *AGO1* and *DCL1* in the moss 14) patens [10, 49], while miR11707 targets *AGO1* in the liverwort, *M. polymorpha* [35]. As for ferns, we found that pmi-miR168 has the potential to target *L. japonicum AGO1* in a site that, given its position in the mRNA, seems to be homologous to the *AGO1* site targeted by miR168 in angiosperms. Since miR168 originated in the lineage leading to ferns and seed plants (Fig 4), it can be hypothesized that the negative feedback loop involving *AGO1* and miR168 has been present throughout the evolutionary history of euphyllophytes. The confirmation of the miR168/*AGO1* regulatory relationship, as well as the study of miR162 and miR403 targets, constitutes an interesting avenue of research in fern regulatory RNA research.

Target prediction of *P. minima* miRNAs yielded two candidate targets that seem to be conserved only between ferns and liverworts. One is the gene encoding polyphenol oxydase (*PPO*), which originated in the early stages of the colonisation of land by plants and has roles in pathogen defence [50]. In both *L. japonicum* and the liverwort, *P. endiviifolia*, *PPO* is targeted at the 3’-UTR by miR408. Another common potential target is RKD, a gene encoding a transcription factor of the RWP-RK family. In the liverworts, *P. endiviifolia* and *M. polymorpha*, *RKD* is targeted at the 3’-UTR by miR319 as revealed by degradome analyses [34, 35]. Similarly, we observed that a *L. japonicum* homologue of RKD is predicted to be targeted at the 3’-UTR by pmi-miR319 and pmi-miR159. Recently, it has been found that RKD is necessary for proper germ cell development in liverworts, as RKD-deficient *M. polymorpha* plants exhibit egg cells with aberrant cellular differentiation and proliferation properties, along with defects in gemma cup formation, which are vegetative reproductive organs [51, 52]. Work in *A. thaliana* indicates that *RKD* homologues might have related functions in egg development of seed plants [53]. Interestingly, the overexpression of miR319 in *M. polymorpha* causes defects in thalli growth and gemma cup formation in gametophytes [36]. Thus, given the important functions of *RKD*, its potential regulation by miR319 in liverworts and ferns might represent an ancient gene regulatory interaction that plays a role in gametophyte growth and germ cell differentiation in basal land plants.

Despite their ecological importance and their key phylogenetic position as sister group to seed plants, the genetic and molecular mechanisms underlying fern development and physiology are poorly known. We hope that our work will add to the growing genomic resources available to researchers dedicated to the study of this group of land plants.

## Materials and Methods

### Plant material

Specimens of *Pleopeltis minima* (Bory) J. Prado & R.Y. Hirai (= *Polypodium squalidum* Vell.; = *Pleopeltis squalida* (Vell.) de la Sota.) [20] were collected in Departamento General Alvear, Corrientes Province, Argentina, near the Uruguay river, in mid-2011. Annual precipitation in the area reaches 17,000 mm approximately. No special permits were needed to collect the specimens. The ferns were found growing on branches of large trees, and a couple of specimens with the branches attached were maintained in a greenhouse at 23C under a 16h/8h light/dark cycle.

### RNA preparation and high-throughput sequencing

Two independent samples of *P. minima* fertile fronds from two different specimens each were subjected to a protocol for total RNA extraction enriched for small RNAs using the mirVana™ miRNA Isolation Kit (Thermo Fisher Scientific). The quantity and quality of the total RNAs were analised with a NanoDrop 2000 spectrophotometer (Thermo Fisher Scientific) and an Agilent 2100 Bioanalyzer (for concentration, 28S/18S and RIN detection; Agilent Technologies). Small RNA libraries were generated from both *P. minima* independent RNA samples using the Illumina Truseq^TM^ SmallRNA Preparation kit(Illumina, San Diego, USA). Total sRNAs were ligated to 3p and 5p adapters (ADTs), and the corresponding cDNA was obtained by reverse-transcription PCR. The purified cDNA library was used for cluster generation on Illumina’s Cluster Station and then sequenced on Illumina GAIIx (LC Sciences, Houston, USA). Raw sequencing reads were obtained using Illumina’s Sequencing Control Studio software version 2.8 (SCS v2.8) following real-time sequencing image analysis and base-calling by Illumina’s Real-Time Analysis version 1.8.70 (RTA v1.8.70).

### Identification of conserved miRNAs

Initial analysis of reads were done with the ACGT101-miR bioinformatics program (LC Sciences, Houston, USA). Raw reads were filtered to remove adaptor sequences and reads which were too short (<15 nt) or too long (>40 nt). Next, sequences corresponding to mRNA fragments, rRNA, tRNA, snoRNA and snRNA mapping to Rfam (http://rfam.janelia.org) were removed, as well as reads mapping to RepeatMasker (http://www.repeatmasker.org/). The resulting non-redundant sRNA reads were mapped to plant precursor and mature miRNA sequences stored in miRBase 21.0 (http://www.mirbase.org/; Kozomara and Griffiths-Jones, 2014) using BLASTN software to identify conserved miRNAs. Length variation (1-3 nt) at both 3’ and 5’ ends and up to two mismatches inside of the sequence were allowed in the alignment. The identified *P. minima* miRNAs were compared to a recent re-evaluation of miRBase entries [8] to discard low-confidence miRNAs from our sample.

To better characterize the phylogenetic distribution of plant miRNAs, *P. minima* sequences were also checked against a large plant miRNA database generated recently [19] which includes the aquatic fern, *M. quadrifolia*. The miRNAs that were reported for *M. quadrifolia* by Chávez-Montes et al [19] were reanalysed more stringently in the following way: First, miRNA sequences for each putative miRNA family were compared against miRBase (http://www.mirbase.org/) and miRNA sequences that exhibited low similarily to miRBase entries (E-value of less than 0.01) were discarded. Next, the abundance of each miRNA sequence was checked against the Comparative Sequencing of Plant miRNA database (http://smallrna.danforthcenter.org/) [19] and sequences that were expressed at low levels (less than 1 read per million) were discarded. After this conservative reassessment, several miRNAs that seemed to be present in *M. quadrifolia* and other plants [19] were discarded from our evolutionary analysis. The resulting bona fide conserved miRNAs were incorporated into the descriptions of miRNA distribution and evolution shown in Figs 3 and 4.

### miRNA target prediction

The P. minima conserved miRNA sequences identified (57 in total; Table 1) were used to search against the transcriptome of the fern, *Lygodium japonicum* [17],for potentialtargets using the psRNATarget platform (http://plantgrn.noble.org/psRNATarget/) [39]. Searches were performed against the Oases_K49 database (downloaded from http://bioinf.mind.meiji.ac.jp/kanikusa/) that consists of a transcriptome assembled from sequenced RNA samples from prothalli, trophophylls, rhizomes and sporophylls obtained using Roche 454 GSFLX and Illumina HiSeq sequencers [17]. Search parameters were Expectation (E) of ≤ 3.0 (which measures sequence complementarity) and UPE of ≤ 25.0 (which measures target accessibility) [39]. For some miRNAs, namely pmi-miR168, the E parameter was relaxed (≤ 4.0) to identify further potential targets.

### Data availability

The raw sequencing data of *P. minima* small RNA transcriptome was submitted to the NCBI Sequence Read Archive under the accession number SRR4294182.

## Acknowledgments

The authors thank Dr. Sara Maldonado (University of Buenos Aires) for critically reading the manuscript.

## Author Contributions

Conceived and supervised research: AMZ. Performed the experiments: FB PIP LMT. Analysed the data: FSJS SIN AMZ. Contributed reagents/materials/analysis tools: MR ANM. Funding adquisition: ANM AMZ. Wrote the paper: AMZ.

## Supporting Information

**S1 Fig. Predicted targeting of fern C3HDZ mRNAs by miR166. (A)** Sequence of a transcript (Locus_1230) encoding a class III homeodomain-leucine zipper protein (C3HDZ) homologue from the fern *L. japonicum*. The region predicted to be targeted by pmi-miR166 is indicated in yellow. The coding region is highlighted in blue. **(B-D)** C3HDZ transcripts belonging to three clades (C3HDZ1, C3HDZ2 and C3HDZ3, respectively) from the fern Psilotum nudum [40]. The coding regions and the regions predicted to be targeted by pmi-miR166 are indicated in blue and yellow, respectively.

**S2 Fig. *Lygodium japonicum TAS3* RNAs targeted by miR390. (A)** *TAS3* RNAs (Locus 20755 and Locus 39179) from *L. japonicum* predicted to be targeted by pmi-miR390. The regions predicted to pair with miR390 are indicated in blue, and the trans-acting small-interfering RNAs (tasi-RNAs) derived from *TAS3* processing are shown in yellow. The tasi-RNAs are predicted to target mRNAs encoding ARF3/4 transcription factors. **(B)** *TAS3* RNAs were aligned with the CLUSTAL Omega program. Regions predicted to be targeted by pmi-miR390 are indicated in blue, with the base pairing with miR390 indicated. Note that the pairing between pmi-miR390 is more extensive at the 5’ site than at the 3’ site, indicating that RISC-mediated cleavage of the RNAs probably occurs only at the 5’ site. The trans-acting small-interfering RNAs (tasi-RNAs) derived from *TAS3* processing are shown in yellow. The tasi-RNAs are predicted to target mRNAs encoding ARF3/4 transcription factors. **(C)** Sequence of a mRNA (isotig 23398) encoding a ARF3/4 transcription factor from *L. japonicum* predicted to be targeted by tasi-RNAs. The coding region is shadowed grey. The segment highlighted in yellow has almost perfect base-pair complementarity with the tasi-RNAs derived from the *TAS3* RNAs shown above.

**S3 Fig. *Lygodium japonicum* RWP-RK mRNA targeted by Pleopeltis miR159 and miR319.** Sequence of a mRNA encoding a RWP-RK transcription factor (Locus 3676) from *L. japonicum* predicted to be targeted by pmi-miR408. The targeted region is located at the 3’-UTR of the mRNA (yellow). The coding region is highlighted in blue. This putative fern miR159/319 target is homologous to an experimental miR319 target of the liverworts, *P. endiviifolia* and *M. polymorpha*.

**S4 Fig. Predicted targeting of fern *PPO* mRNA by miR408.** Sequence of a mRNA encoding Polyphenol oxydase (PPO, Locus 5437) from *L. japonicum* predicted to be targeted by pmi-miR408. The targeted region is located at the 3’-UTR of the mRNA (yellow). The coding region is highlighted in blue. This putative fern miR408 target is homologous to an experimental miR408 target of the liverwort, *P. endiviifolia*.

**S5 Fig. Predicted targeting of fern AGO1 mRNA by miR168. (A)** Sequence of a transcript (Isotig21625) encoding an Argonaute 1 (AGO1) homologue from the fern *L. japonicum*. The region predicted to be targeted by pmi-miR168 is indicated in yellow. The coding region is highlighted in blue. **(B)** Comparison between the experimentally described pairing between ath-miR168a and AGO1 of A. thaliana and the predicted pairing between pmi-miR168_v1 and the AGO1 mRNA of *L. japonicum*. The AGO1 sequences recognized by miR168 are highlighted in yellow. Codons are indicated in alternating red and black lettering and the encoded aminoacid residues are shown below the codons.

**S1 Table. Statistics of sRNA libraries.**

**S2 Table. *P minima* putative 5p and 3p miRNAs duplexes.**

